# isoformant: A visual toolkit for reference-free long-read isoform analysis at single-read resolution

**DOI:** 10.1101/2021.12.17.457386

**Authors:** Daniel D. Le, William Stephenson, Faye T. Orcales

## Abstract

`isoformant` is an analytical toolkit for isoform characterization of Oxford Nanopore Technologies (ONT) long-transcript sequencing data (*i*.*e*. direct RNA and cDNA). Deployment of these tools using `Jupyter Notebook` enables interactive analysis of user-defined region-of-interest (ROI), typically a gene. The core module of `isoformant` clusters sequencing reads by *k*-mer density to generate isoform consensus sequences without the requirement for a reference genome or prior annotations. The inclusion of differential isoform usage hypothesis testing based on read distribution among clusters enables comparison across multiple samples. Here, as proof-of-principle, we demonstrate the utility of `isoformant` for analyzing isoform diversity of commercially-available isoform standard mixtures. `isoformant` is available here: https://github.com/danledinh/isoformant.

## Introduction

To capture the full potential of nanopore-based transcriptome analysis, novel computational approaches are needed to extract information beyond gene identity, such as RNA base modifications and isoform usage.^1^ Conceptually, long-read sequencing enables direct observation of exon/intron arrangement along an isoform. However, in practice, relatively high sequencing error rate and truncations (*e*.*g*. non-processive reverse transcription, RNA degradation) cause alignment artifacts that complicate isoform characterization. Authors of `FLAIR`^2^ attempt to address these challenges by correcting splice junction sequences using prior splice site annotations. `StringTie2`^3^ is another approach based on an assembly process, which does not require supplemental annotations to correct splice junction sequences. Benchmarking by the authors of `StringTie2` suggests that their approach is comparable to or outperforms `FLAIR` across several sample types. Here, we present `isoformant` as an alternative approach that derives isoforms by generating consensus sequences from long reads clustered on *k*-mer density without the requirement for a reference genome or prior annotations. In principle, sequence homology based on *k*-mer spectra of noisy long reads is tolerant of errors^4^, especially when compared to base-resolution alignment strategies, which makes it suitable for nanopore-based isoform analysis. This analytical toolkit allows users to interactively and visually explore isoform data at the granularity of individual reads.

`isoformant` was developed based on the concept that an individual long-read isoform can be uniquely identified by its constituent *k*-mer composition. Namely, for an appropriate length *k*, each unique read in a mixture can be represented by a correspondingly unique *k*-mer frequency vector. Thus, a mixture of *m* long reads can be represented by a *m* x *n* matrix, where *n* = 4^*k*^. The process by which `isoformant` extracts isoform information from such matrices was inspired by the analytical pipeline outlined by the single-cell transcriptomics package `SCANPY`.,^5,6^ Beyond convenient implementation of analytical tools, `SCANPY` provides a scalable framework for handling large matrices and associated metadata. Moreover, `isoformant` provides tools for visualizing isoform sequence variation and performing differential isoform usage hypothesis testing. The toolkit is intended for interactive analysis using `Jupyter Notebook`.

## Methods

### isoformant toolkit

Detailed documentation can be found here: https://isoformant.readthedocs.io/en/latest/isoformant.html. Briefly, the `isoformant` toolkit is composed of the following core modules:

0. Read alignment (prior to `isoformant` usage) Minimap2^7^ (version 2.21) with `-ax splice` flag is recommended to align reads.
1. Preprocessing reads One or more coordinate-sorted and indexed `.bam` alignment file paths (`.bai` index file must be in same directory) can be specified for processing. Reads are restricted to a user-defined ROI, typically a gene. Then reads are filtered by mean base quality (default = 10) and length (default = 300). If more than one. `bam` file, post-filter read depth is balanced by random downsampling to the .`bam` file with fewest filtered reads. In addition, the user can specify the maximum number of passing reads per .`bam` file (default = 1000).
2. *k*-merization and clustering For each passing read, the *k*-mer density (*k*-mer frequency, computed using the `khmer` software^8^, divided by total number of observed *k*-mers; default *k* = 7) is computed and appended to a matrix *M*. This matrix along with collated metadata are combined in an `Anndata` object. All subsequent transformations are performed on this object. First, dimensionality reduction was performed using Principal Component Analysis (PCA) of the *k*-mer density matrix *M*, which yields a Principal Component (PC) matrix *M*\* (default = 100 PCs). Second, the connectivity among reads is computed for a *k*-nearest neighbor (KNN) graph based on *M*\*. The resultant KNN graph is the basis for visualization via Uniform Manifold And Projection (UMAP)^9^ and for Leiden clustering^10^ to identify putative isoforms.
3. Consensus calling For each cluster, a random sample of reads (default = 10) is selected for consensus calling using the Partially-Ordered Alignment (POA) algorithm.^11^ POA was designed to handle gapped multiple alignments, which is necessary to determine alternative splice variants.
4. Hypothesis testing In cases where multiple samples are simultaneously analyzed, a chi-squared test can be used to determine differential isoform usage among the samples. The test assumes balanced read depth among samples (see “1. Preprocessing reads”); therefore, the null hypothesis assumes uniform sample representation in each cluster (*i*.*e*. putative isoform). For example, given sample *A* and sample *B*, it is expected that an isoform cluster is comprised of an equal number of reads from both samples. For any given cluster, rejection of the null hypothesis indicates significant imbalance in sample representation. In such cases, differential isoform usage among samples is determined at the level of individual clusters. Because the test is performed on all clusters, Bonferroni multiple-comparison correction is applied, yielding adjusted P values.

### SIRV cDNA sequencing

The SIRV cDNA library for nanopore sequencing was generated according to the 10X Genomics based single cell cDNA library preparation from Lebrigand *et al*. 2020 which depletes cDNA that lacks poly(A)/poly(T) sequences.^12^ Briefly, ∼5 ng SIRV Set 4 (Lexogen) was reverse transcribed (10X Genomics Gel Bead Primer - polydT) with template switching (10X Genomics Template Switch Oligo). Following RT, PCR (5 cycles) was performed with a 5’ biotinylated forward primer and non-biotinylated reverse primer. After 0.6X SPRIselect purification to remove excess biotinylated primers, biotinylated cDNA was bound to Dynabeads M-270 Streptavidin beads (Thermo Fisher Scientific). After washing, on-bead PCR was performed (7 cycles) with non-biotinylated forward and reverse primers with identical sequences as those used in the initial biotinylated amplification step. 110 ng of amplified SIRV cDNA was input for library preparation for nanopore sequencing using the direct cDNA kit (SQK-DC109) proceeding directly from end-prep. Nanopore sequencing produced ∼14 M reads with median read length and PHRED quality score of 1170 bases and 12.3, respectively.

## Results

To demonstrate the features of `isoformant`, we analyzed publicly-available direct RNA sequencing data of Spike-In RNA Variant (SIRV) control mixes, generated from an ONT Minion device.^13^ Specifically, two SIRV-Set 1 isoform mixes were considered: 1) The E0 mix contains equimolar amounts of each isoform (NCBI accession: SRX3204588), and 2) The E2 mix contains variable known amounts of each isoform, which allows for differential usage analysis (NCBI accession: SRX3204589). Here, we report the results for an ROI spanning the SIRV1 reference sequence, composed of 8 annotated isoforms.^14^ Minimap2 was used for aligning reads to the SIRV reference sequences.^15^ The SIRV E0 and E2 .`bam` files were preprocessed by `isoformant (max_reads = 5000, qual_cutoff = 10, len_cutoff` = 300), yielding 31,024 and 10,494 passing reads, respectively. By default, read depth is balanced among samples and limited to the `max_read` number of reads; therefore, the resultant merged dataset was composed of 5,000 passing reads from each of the two samples. Then, *k*-merization (`ksize` = 7) and clustering (`n_pcs = 50, n_neighbors = 15, min_dist = 0.1, resolution = 1`) were performed, yielding the UMAP in Figure 1.

**Figure 1:**
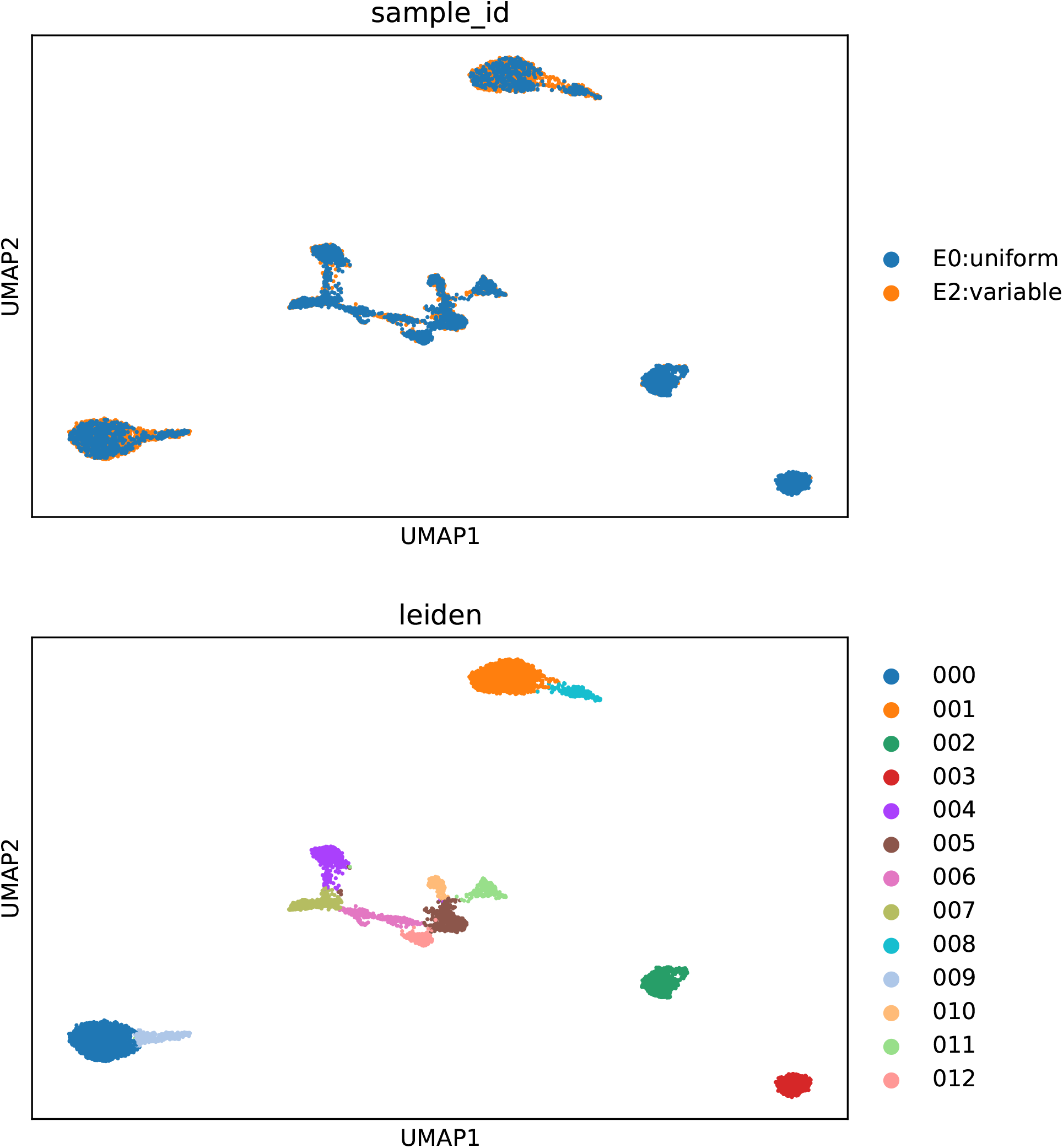
UMAP of reads aligned to SIRV1. (top panel) color-coding by sample. (bottom panel) color-coding by Leiden cluster.

Next, randomly-sampled reads (`bam_n = 20`) from each Leiden cluster in Figure 1 were used for consensus calling. The resultant sequences were aligned to the SIRV reference sequences^15^, which yielded the cluster-specific alignment tracks in Figure 2. Some cluster-specific consensus alignment tracks were matched to reference tracks^14^ based on sequence similarity in terms of shortest Levenshtein distance between pairs, yielding the following cluster:reference assignments: (010:SIRV101), (012:SIRV102), (005:SIRV103), (000:SIRV105), (004:SIRV106), (001:SIRV107), (002:SIRV108), and (003:SIRV109). Together with the UMAP representation of reads sequence homology, it was possible to infer relatedness among read clusters. For example, the cluster 000 (*i*.*e*. SIRV105) is fused with unassigned cluster 009, which contains reads that lack aligned 5’ exons. Thus, cluster 009 reads are likely truncation products or are misaligned due to relatively high base calling error rate.

**Figure 2:**
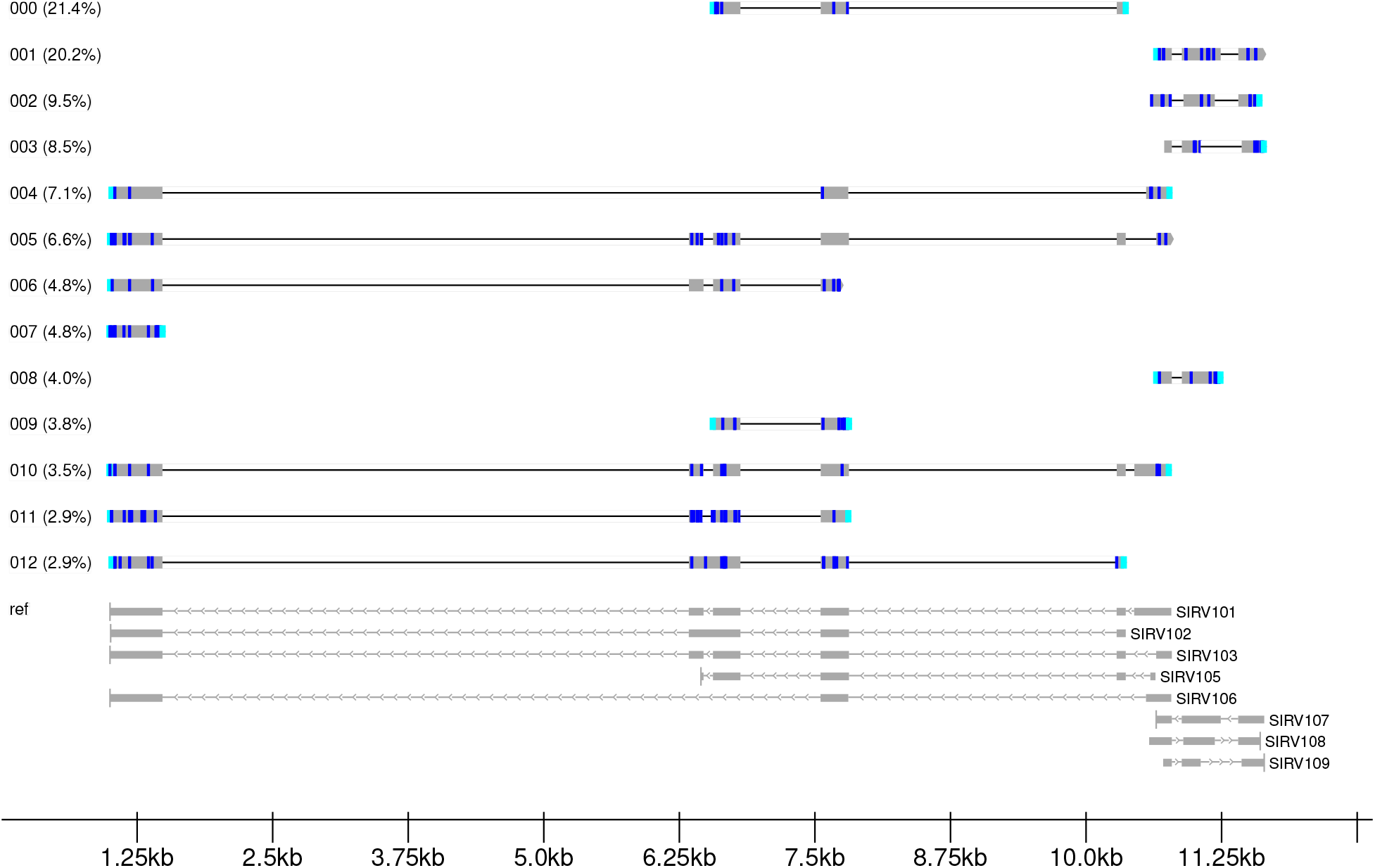
Cluster-specific consensus alignment tracks. (color code) substitution:blue, N-ambiguous:yellow, insertion:red, deletion:green, clipping:cyan.

To determine differential isoform usage between the E0 and E2 isoform mixes, a chi-squared test was performed for each read cluster. Because sample read depth was balanced, the null hypothesis assumes equal probability of observing reads originating from either E0 or E2 libraries. For clusters representing isoforms, rejection of the null hypothesis indicates differential isoform usage. For the remaining clusters originating from truncation or sequencing error, among other sources of variation, the chi-squared test identifies differential abundance between E0 and E2 libraries. Of the clusters identified as SIRV sequences, cluster 003 (*i*.*e*. SIRV109) showed the most significant (adj. P value = 5.52684e-169; see Table 1) differential isoform usage between the E0 and E2 mixes. This is consistent with the fact that SIRV109 abundance in E0 mix is 32-fold higher than in E2 mix, the highest disparity among the SIRV1 isoforms, according to product specifications. The observed relative abundance of SIRV109 in cluster 003 between E0 and E2 mixes is approximately 42-fold (or about 31% over-estimated), shown in Figure 3. Extension of this analysis to all assigned clusters, displayed in Figure 4, reveals strong correlation between observed and expected SIRV1 isoform proportions (Pearson’s *r*^*2*^ = 0.97). Moreover, differential isoform usage analysis was performed on a biological sample: the Universal Human Reference (UHR) transcriptome.^16^ Genome-wide differential isoform usage analysis of direct mRNA sequencing reads from UHR (NCBI accession: SRR12010483) was performed using a recently developed tool called `LIQA`.^17^ Among the genes with deepest coverage that exhibited differential isoform usage was SNRPB. `LIQA` estimated 36% and 63% relative abundance of two transcript isoforms: Gencode ENST00000438552.6 (short 3’-exon) and ENST00000381342.7 (long 3’-exon), respectively. Whereas, `isoformant` estimated the relative abundances to be 43% and 57% (excluding an observed truncation population; see Figure 5) a discrepancy relative to `LIQA` of approximately 6-7%. Taken together, these results demonstrate that `isoformant` accurately quantifies isoform abundances; thus, enabling the detection of differential isoform usage both within and across samples.

**Table 1:** Chi-squared hypothesis test results comparing read frequency in each cluster between E0 and E2 libraries. Bonferroni correction was used to adjust P value for multiple comparisons.

| leiden | P value | adj. P value |
| --- | --- | --- |
| 003 | 4.25142e-170 | 5.52684e-169 |
| 001 | 4.66971e-156 | 6.07063e-155 |
| 002 | 2.3521e-96 | 3.05773e-95 |
| 000 | 2.29408e-74 | 2.9823e-73 |
| 012 | 2.58124e-37 | 3.35561e-36 |
| 008 | 9.61311e-22 | 1.2497e-20 |
| 009 | 3.15417e-16 | 4.10042e-15 |
| 006 | 1.09875e-14 | 1.42838e-13 |
| 004 | 1.92015e-14 | 2.49619e-13 |
| 007 | 3.67724e-08 | 4.78041e-07 |
| 005 | 7.89757e-08 | 1.02668e-06 |
| 011 | 0.000344158 | 0.00447405 |
| 010 | 0.215613 | 1 |

**Figure 3:**
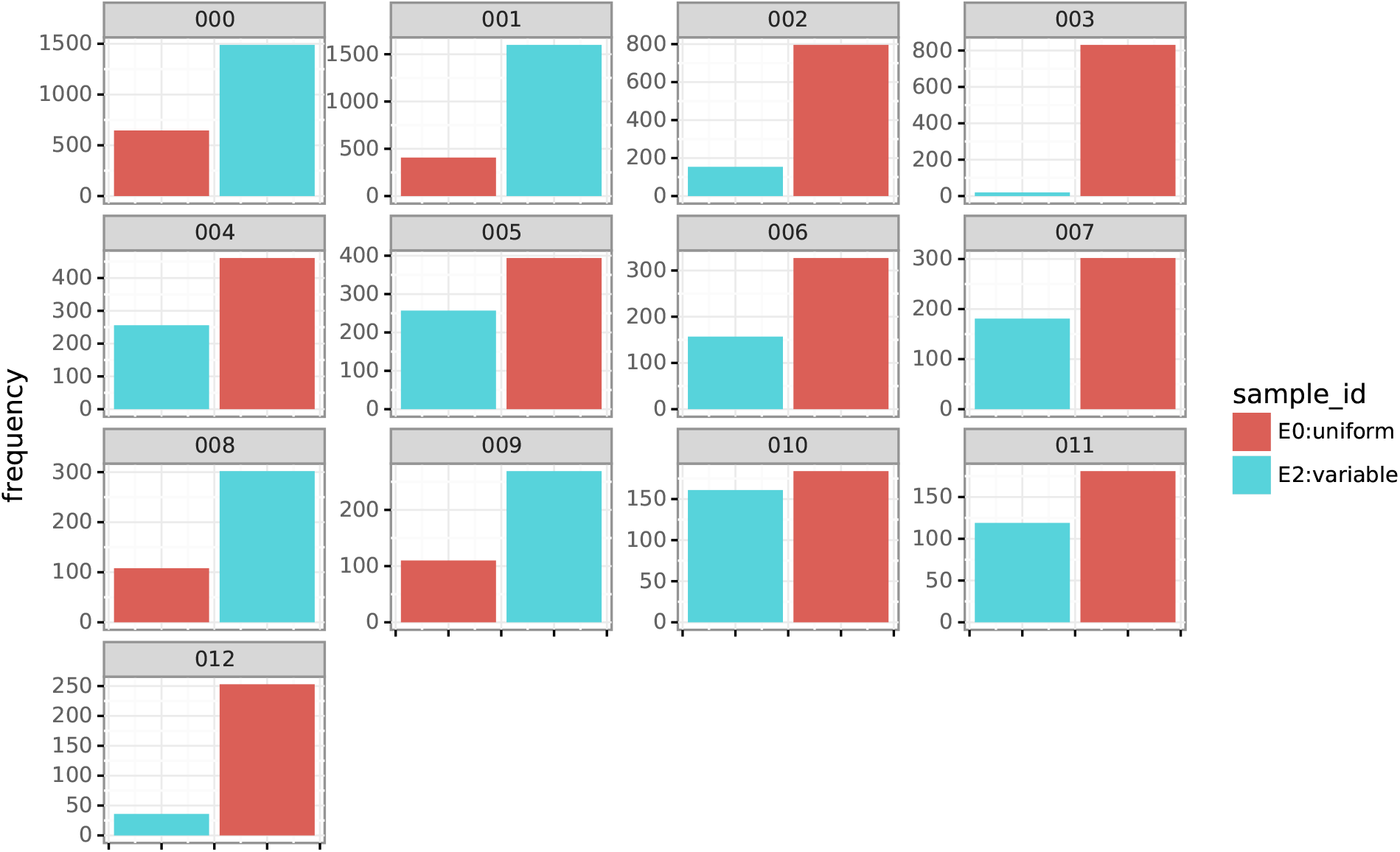
Relative read abundance between E0 and E2 libraries among clusters.

**Figure 4:**
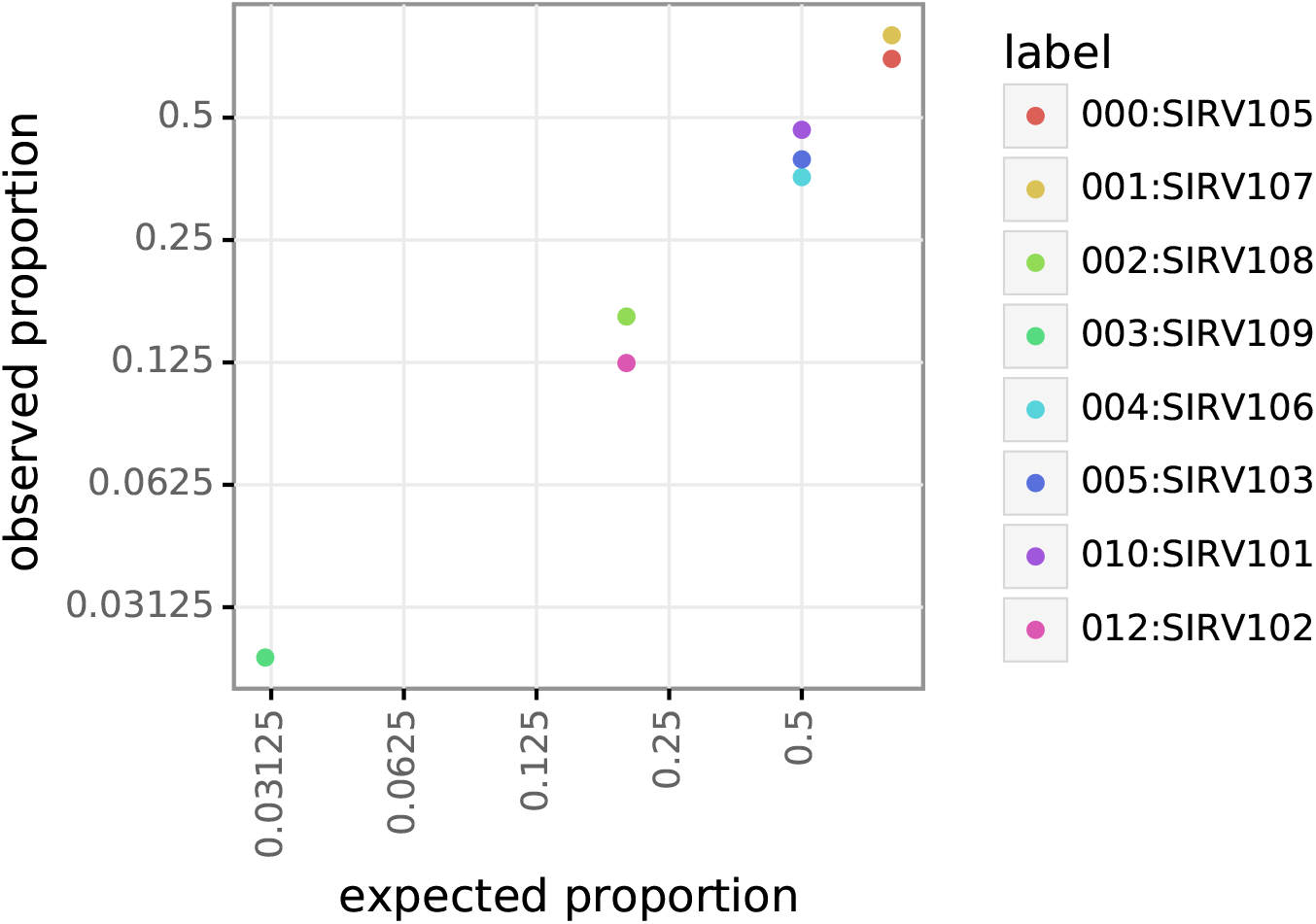
Correlation of expected and observed proportions of E0 and E2 reads by assigned cluster.

**Figure 5:**
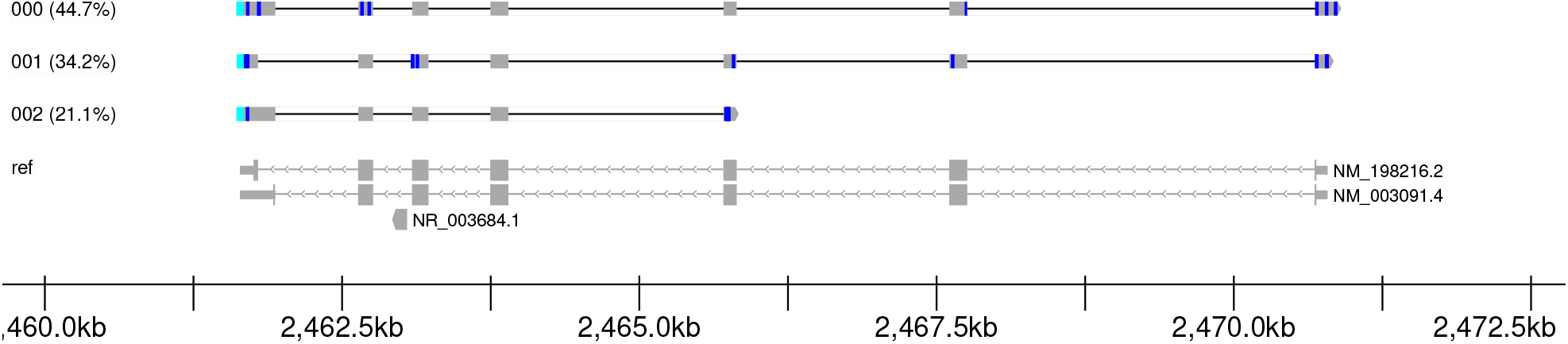
SNRPB isoform alignment tracks and relative abundances from analysis of the Universal Human Reference. (color code) substitution:blue, N-ambiguous:yellow, insertion:red, deletion:green, clipping:cyan.

Next, `isoformant` was used to compare reads originating from either direct RNA or cDNA sequencing. Given the reported differences in error rate and type (*i*.*e. k*-mer frequency distribution biases) between these sequencing modalities,^18,19,20^ we aimed to characterize how these disparities manifest when using `isoformant` for isoform analysis. Direct RNA reads (as described above) and cDNA reads (see Methods) from the E0 isoform mix were used as inputs to `isoformant`. The analysis was performed using nearly identical parameters as listed above. The only exception was the `resolution` parameter (*i*.*e*. clustering resolution) was reduced to 0.20, which decreased oversegmentation of read groups. From the UMAP embedding of reads aligned to SIRV1, we observed complete segregation of reads based on sequencing modality (see Figure 6). For the cDNA library, reads were clustered into 8 distinct groups corresponding to annotated isoforms (see Figure 7): (011:SIRV101), (002:SIRV102), (003:SIRV103), (006:SIRV105), (007:SIRV106), (009:SIRV107), (013:SIRV108), and (012:SIRV109). Whereas, the direct RNA library exhibited only 4 distinct annotated isoform groups : (008:SIRV105), (010:SIRV107), (005:SIRV108), and (004:SIRV109). In addition, there was 1 fused group consisting of clusters 000 and 001

**Figure 6:**
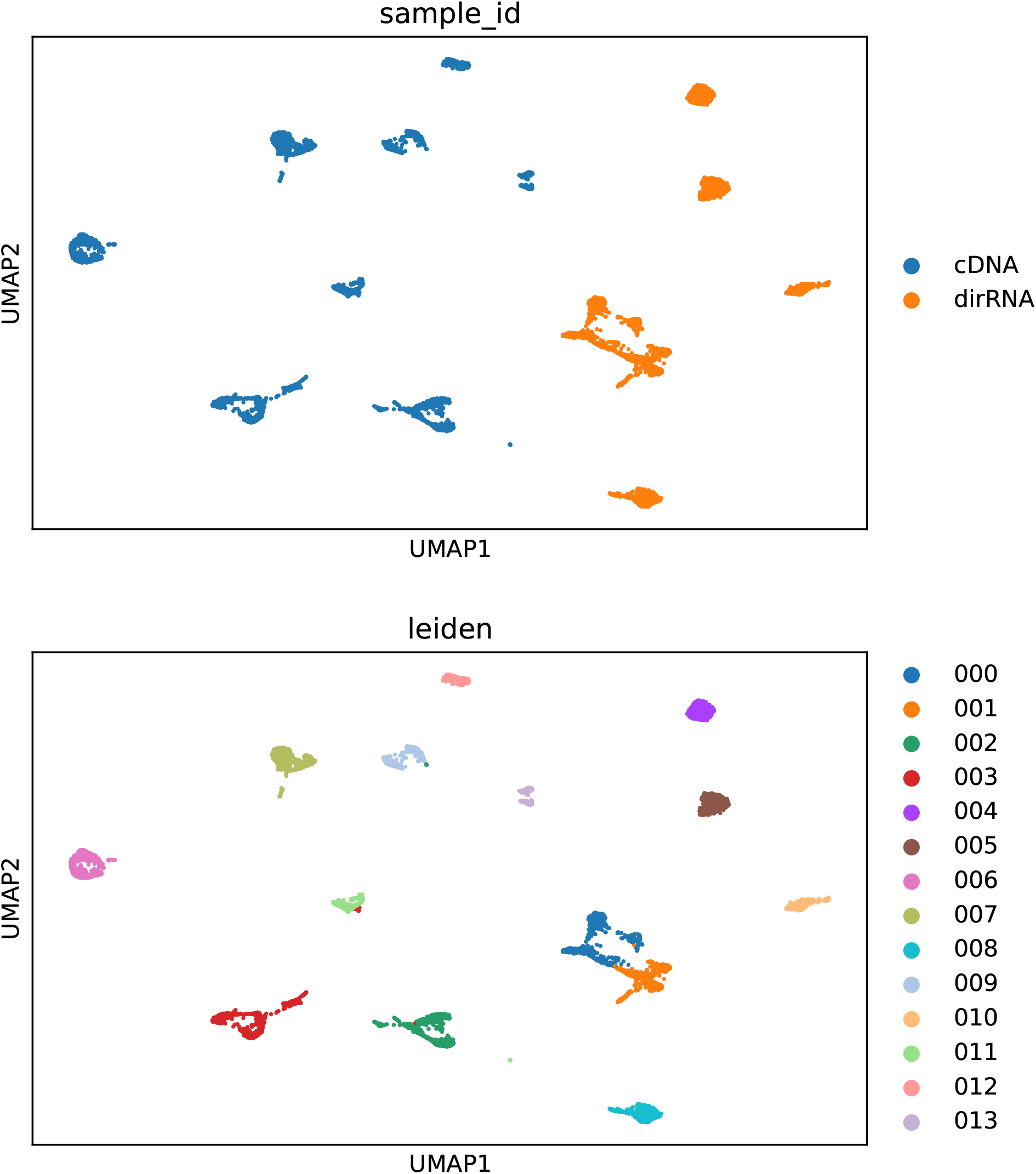
UMAP of direct RNA and cDNA reads aligned to SIRV1. (top panel) color-coding by sample. (bottom panel) color-coding by Leiden cluster.

**Figure 7:**
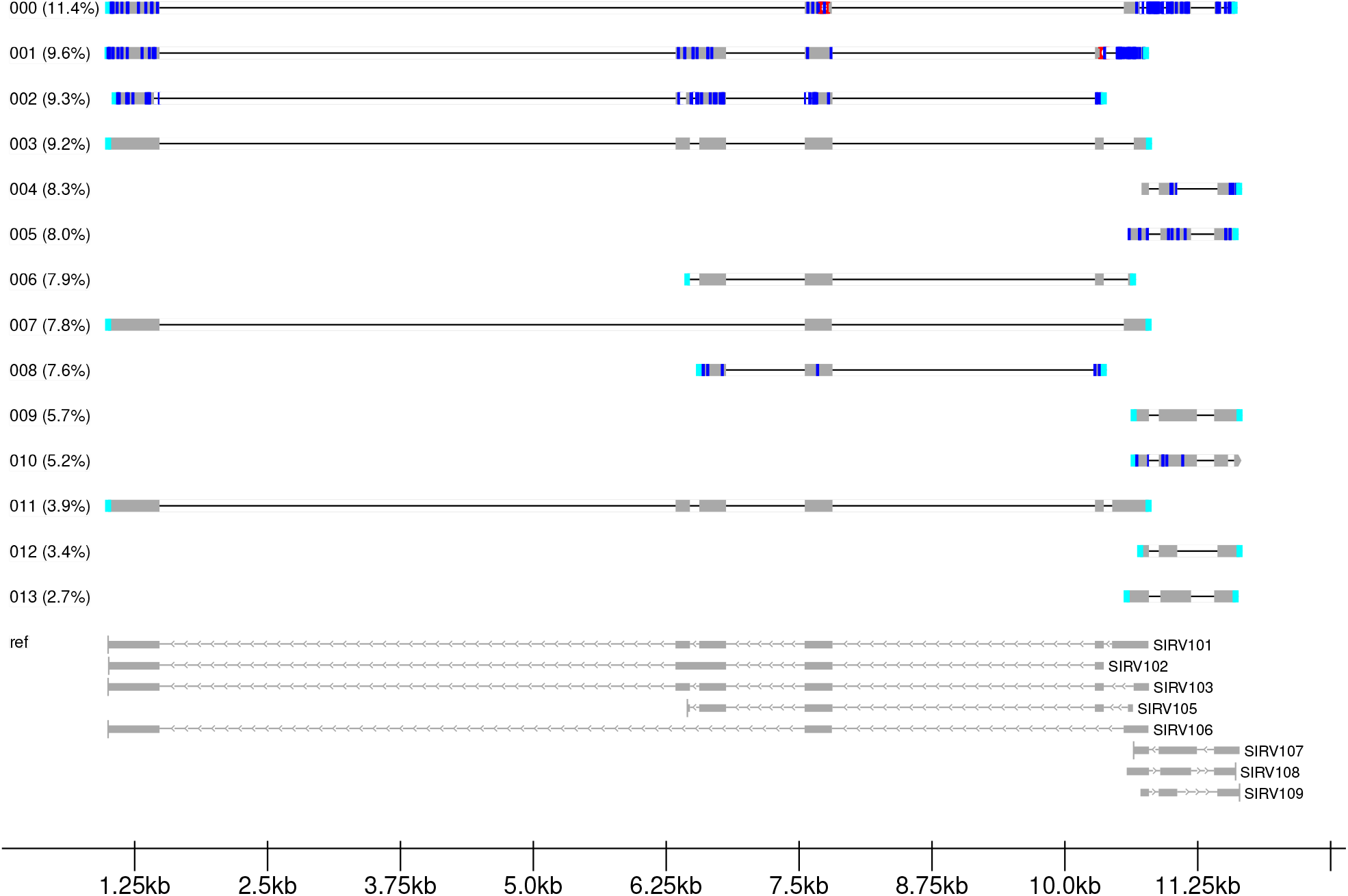
Cluster-specific consensus alignment tracks from direct RNA and cDNA libraries. (color code) substitution:blue, N-ambiguous:yellow, insertion:red, deletion:green, clipping:cyan.

Repeating the pipeline (`res` = 0.35) using only reads from the fused group resulted in a UMAP embedding focused on the sequence variation among this subset of reads (see Figure 8). Leiden clustering yielded 7 discrete read groups, corresponding to the remaining 4 annotated isoforms not discovered in the initial analysis (see Figure 9): (004:SIRV101), (003:SIRV102), (002:SIRV103), and (001:SIRV106). The 3 unassigned clusters were composed of putative artifacts (*e*.*g*. truncation, sequencing error, *etc*.).

**Figure 8:**
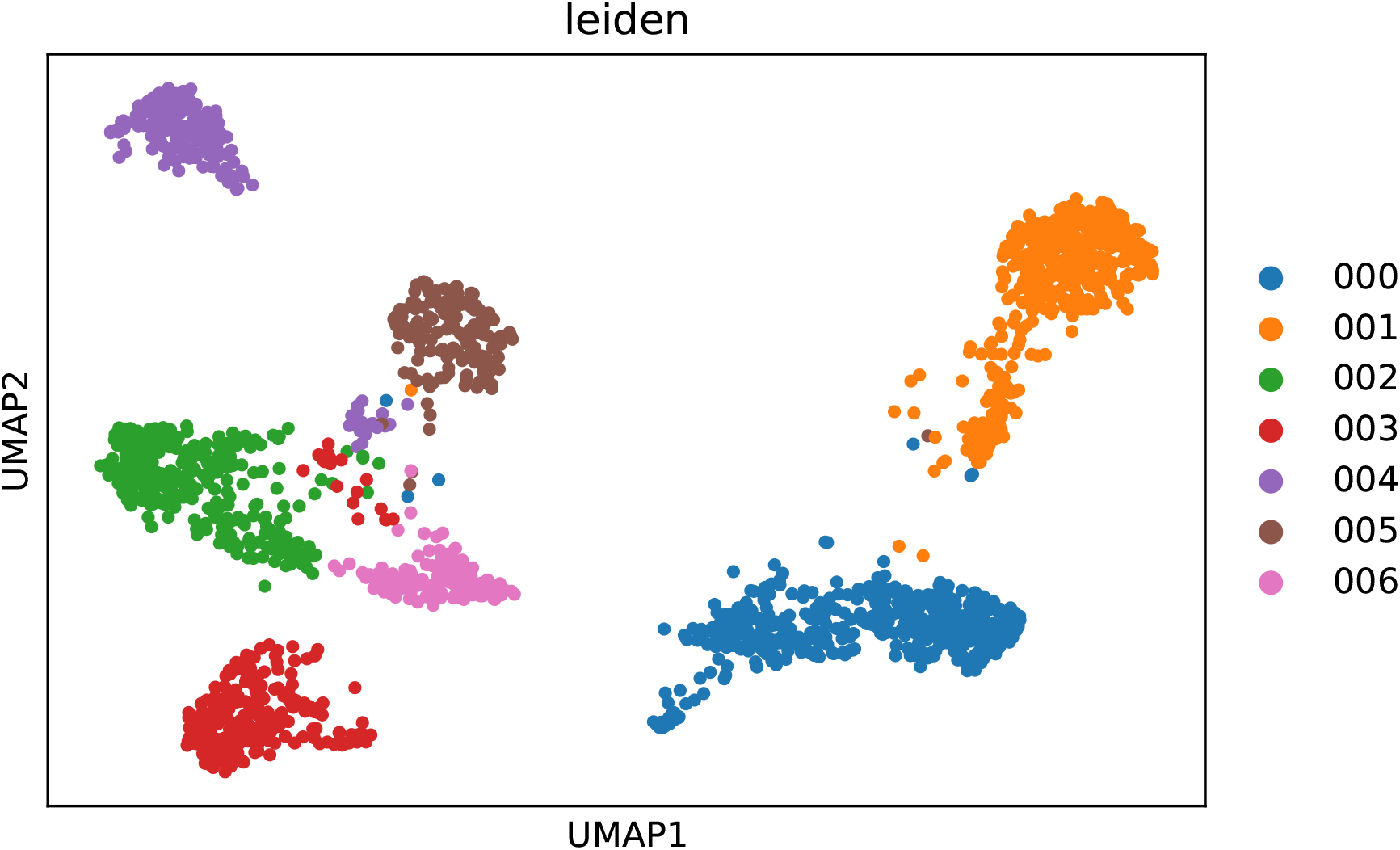
UMAP of direct RNA reads from fused group. Color-coding by Leiden cluster.

**Figure 9:**
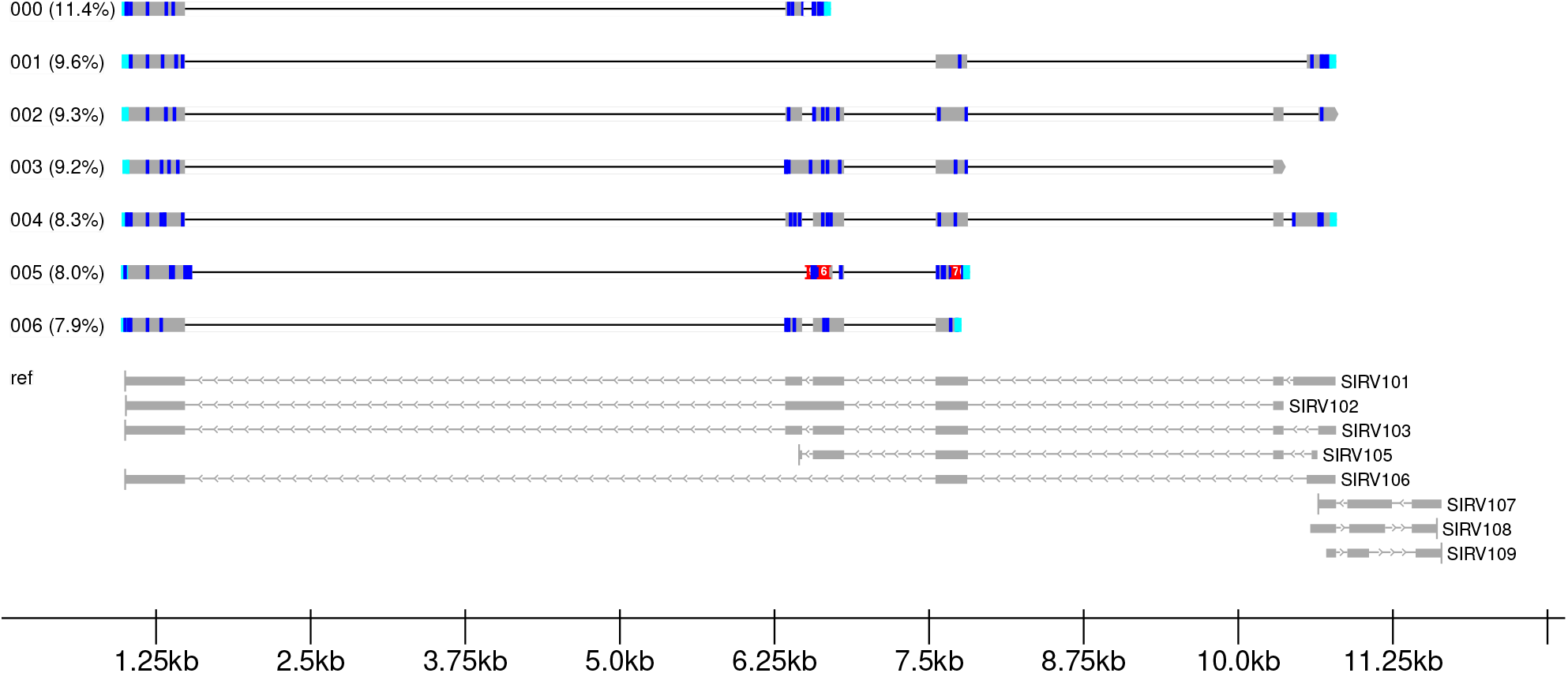
Cluster-specific consensus alignment tracks from fused group reclustering. (color code) substitution:blue, N-ambiguous:yellow, insertion:red, deletion:green, clipping:cyan.

## Discussion

`isoformant` was designed to focus on ROIs; thus, it is not suitable for multi-loci or genome-wide applications. In those cases, `isoformant` may be a useful complement that enables detailed isoform analysis of high-priority loci. It is also worth reiterating the bias associated with sequencing modality (*e*.*g*. direct RNA versus cDNA) on *k*-mer distribution and error rate that may explain some variation in read clustering. In addition, it is plausible that differences in flowcell or basecaller technology may also introduce bias. Another potential limitation of `isoformant` is high memory usage in generating *k*-mer frequency tables when *k* size is large; however, there are several alternate approaches with more efficient memory utilization based on benchmarking.^21,22^ Moreover, rather than precisely counting *k*-mers, scalable and performant approaches such as minhashing with locality sensitive hashing (LSH) approximate sequence similarity among reads.^23,24^ Lastly, `isoformant` currently lacks an isoform classification feature, which limits streamlined comparison to prior annotations. In future versions of `isoformant`, we plan to improve consensus calling by implementing options for polishing using Racon^25^ and for transcript assembly using StringTie2.^3^ In addition, normalization or weighting procedures may be implemented to improve sequence variation resolution. For example, Term Frequency-Inverse Document Frequency (TF-IDF) transformation^26^ can be used to prioritize rare *k*-mers that may better explain the sequence variation among reads.

In this manuscript, we demonstrated the capabilities of the `isoformant` toolkit to aid in long-read isoform characterization. Unlike other tools that output genome-wide isoform annotations, `isoformant` was designed to enable exploration of a ROI in detail. This visual toolkit allows users to inspect individual reads in relation to others, which provides a broader understanding of the isoform diversity landscape. Through the analysis of SIRV control mixes, we demonstrated the ability of `isoformant` to aggregate reads by isoform identity, without the need for splice-junction annotations, and to estimate differential isoform usage. Moreover, we explored how sequencing modality can influence isoform identity determination. Namely, we observed more discrete reads groups matching annotated isoforms in a cDNA library compared to a sample-matched direct RNA library, which exhibited poor segregation among related reads due to basecalling errors. To resolve such ambiguity, we showed that iterative clustering can be successfully employed to resolve the subtle sequence variation among read cluster subpopulations. Taken together, `isoformant` empowers users with fundamental analytical tools to interactively explore long-read isoform data.

